# Interplay of condensate material properties and chromatin heterogeneity governs nuclear condensate ripening

**DOI:** 10.1101/2024.05.07.593010

**Authors:** Deb Sankar Banerjee, Tafadzwa Chigumira, Rachel M. Lackner, Josiah C. Kratz, David M. Chenoweth, Shiladitya Banerjee, Huaiying Zhang

**Affiliations:** Department of Physics, Carnegie Mellon University, Pittsburgh, PA 15213, USA; James Franck Institute, University of Chicago, Chicago, IL 60637, USA; Department of Chemical Engineering, Carnegie Mellon University, Pittsburgh, PA 15213, USA; Department of Chemistry, University of Pennsylvania, Philadelphia, PA 19104; Department of Biological Sciences, Carnegie Mellon University, Pittsburgh, PA 15213, USA; Computational Biology Department, Carnegie Mellon University, Pittsburgh, PA 15213, USA

**Keywords:** Biomolecular condensates, Ostwald ripening, Elastic ripening, Nuclear bodies, Chromatin

## Abstract

Nuclear condensates play many important roles in chromatin functions, but how cells regulate their nucleation and growth within the complex nuclear environment is not well understood. Here, we report how condensate properties and chromatin mechanics dictate condensate growth dynamics in the nucleus. We induced condensates with distinct properties using different proteins in human cell nuclei and monitored their growth. We revealed two key physical mechanisms that underlie droplet growth: diffusion-driven or ripening-dominated growth. To explain the experimental observations, we developed a quantitative theory that uncovers the mechanical role of chromatin and condensate material properties in regulating condensate growth in a heterogeneous environment. By fitting our theory to experimental data, we find that condensate surface tension is critical in determining whether condensates undergo elastic or Ostwald ripening. Our model also predicts that chromatin heterogeneity can influence condensate nucleation and growth, which we validated by experimentally perturbing the chromatin organization and controlling condensate nucleation. By combining quantitative experimentation with theoretical modeling, our work elucidates how condensate surface tension and chromatin heterogeneity govern nuclear condensate ripening, implying that cells can control both condensate properties and the chromatin organization to regulate condensate growth in the nucleus.

## Introduction

The human cell nucleus is a complex environment containing various nuclear bodies embedded in or attached to an expansive network of chromatin (1). These nuclear bodies, such as the nucleoli, histone locus body, PML bodies, and DNA-damage foci, play important roles in facilitating chromatin functions such as transcription, replication and DNA repair (2, 3). There is increasing evidence that nuclear bodies are biomolecular condensates formed by phase separation (4–7). Given the intrinsic ability of the liquid-like condensates to coarsen via coalescence and Ostwald ripening (8), how cells regulate these processes to achieve controlled growth at sites of nucleation become an outstanding question. This question is particularly important for nuclear condensates as many of their chromatin associated functions depend on proper localization (9, 10).

Previous works have shown that the stiffness of the chromatin network can inhibit condensate nucleation (11). In addition, chromatin can inhibit condensate coalescence by reducing condensate mobility and prevent Ostwald ripening by suppressing the growth of large condensates (12, 13). These studies suggest that cells can potentially regulate local chromatin stiffness to control condensate growth in the nucleus.

However, the role of the chromatin organization is less clear. Chromatin is not uniformly distributed in the nucleus but known to have a heterogeneous organization where heterochromatin domains are more dense while the euchromatin regions are less dense (14, 15). This variability in density is additionally regulated by cells, effectively controlling access to genes and influencing nuclear body function and location (2, 3). Chromatin organization goes awry in diseased cells, which may contribute to the aberrant condensate landscape (16, 17). Heterogeneity in chromatin density correlates with heterogeneity in mechanical stiffness, but how this mechanical heterogeneity affects condensate nucleation and growth remains to be learned.

In addition to the chromatin, the condensate properties can also be important in controlling the formation and growth of nuclear condensates. Theory and experiments in colloidal systems revealed that liquid droplets can grow in a polymer network if the condensation pressure is larger than the elastic forces from the polymer (18). On the other hand, gradients in stiffness can inhibit Ostwald ripening or drive elastic ripening, depending on the relative strength of stiffness gradient to surface tension (19). This suggests that cells potentially regulate both chromatin stiffness and condensate properties to control condensate growth in the chromatin. Biomolecular condensates are known to have a wide range of material properties such as surface tension and viscosity (20). However, it is not known whether surface tension of nuclear condensates can be in the range to counter the influence of chromatin stiffness and thus can be exploited by cells to fine-tune condensate nucleation, growth, and sizes in the nucleus.

In this work, we address these outstanding questions by using a chemical dimerizer to induce two types of nuclear condensates in different chromatin environments for comparative growth dynamics assessment. We observe that both types of condensates can grow through coalescence, diffusion, and ripening. However, the proportions of each growth mode are different for the different types of condensates. To explain the experimental observations, we developed a physical model for condensate growth in a heterogeneous elastic environment that represent the chromatin network. In particular, we considered the effect of size-dependent mechanical pressures that condensates may experience from the surrounding chromatin. Our model captures the experimentally measured condensate growth dynamics and predicts the stability of condensates based on their surface tension and the surrounding chromatin stiffness. The model predicts that the coarsening dynamics for different condensates are affected differently by changes in the chromatin landscape, which we confirmed experimentally by perturbing the heterogeneity in chromatin organization. Together, our work shows that the interplay between condensate surface tension and chromatin mechanical heterogeneity controls condensate growth in the heterogeneous physical environment of the nucleus. This indicates that cells can regulate both the material properties and the chromatin organization to control condensate growth, and the aberrant nuclear condensate landscape in diseased cells can be attributed to abnormality in condensate composition and chromatin organization.

## Results

### Condensates made with different proteins have different material properties

To test whether condensate material properties affect condensate growth in the nucleus, we selected a coiled-coil protein and a disordered protein to generate condensates that we predicted to have different material properties. For the coiled-coil protein, we chose Mad1, a human mitotic protein that has the propensity to form condensates (21). For the disordered protein, we chose the intrinsically disordered region of the *C. elegans* p-granule protein LAF-1 that is rich in arginine-glycine-glycine (RGG) repeats (22). We used a previously described chemical dimerization system to induce condensates by linking the phase separation proteins to an oligomer in the nucleus of U2OS cells (23–25) (Fig. 1A). The dimerizer Trimethoprim-Fluorobenzamide-Halo ligand (TFH), consisting of chemically linked Trimethoprim and a Halo-ligand that interact with eDHFR and Halo enzyme, respectively, can dimerize proteins fused to eDFHR and Halo (26) (Fig. 1A). Using THF to dimerize LacI, a dimer protein, to Mad1 was enough to initiate the coiled-coil condensate formation (Movie 1). A hexamer, Hotag3 (25), was required to be dimerized to RGG to induce the disordered condensates in the nucleus (Movie 2).

**Fig. 1.**
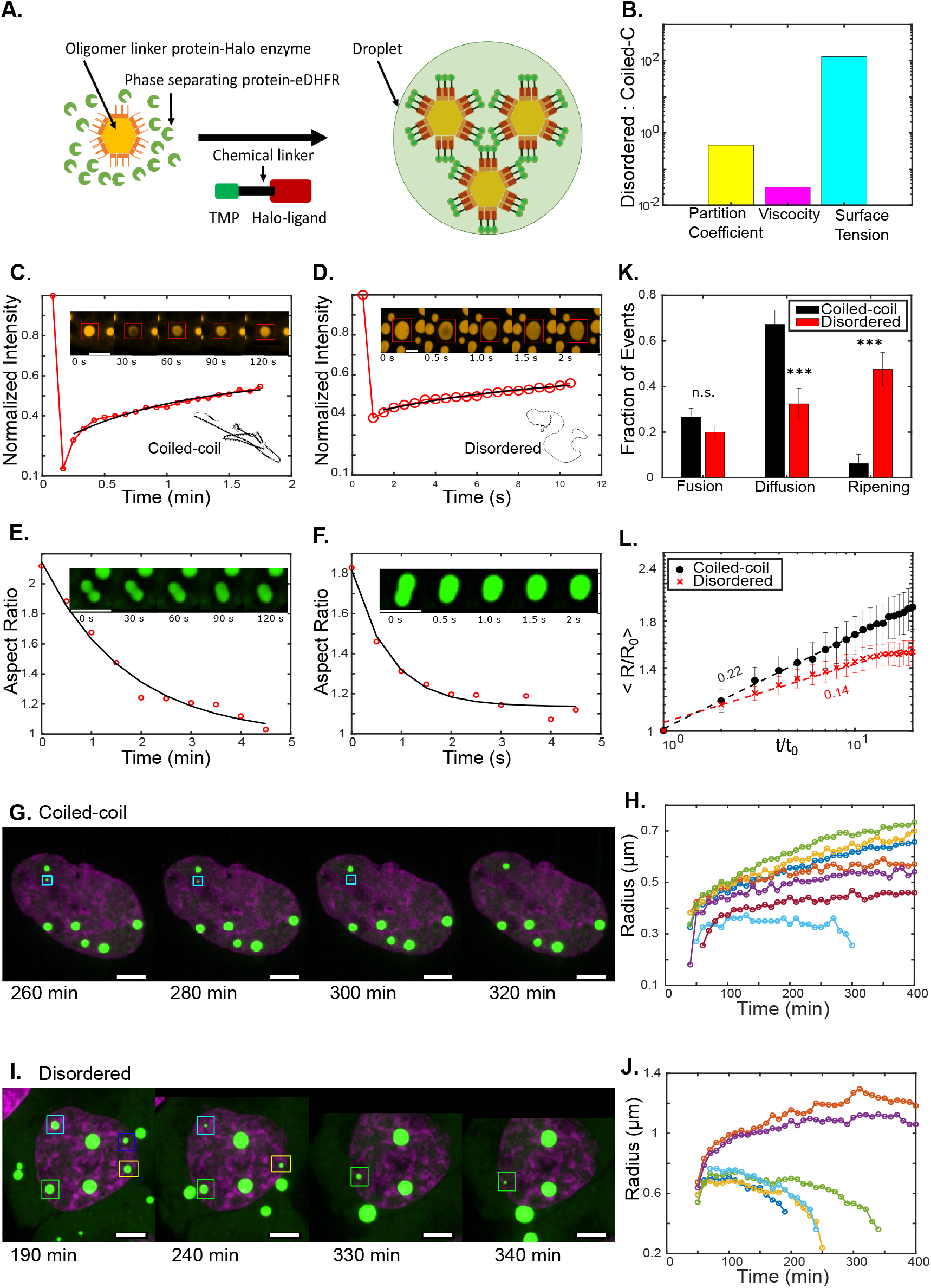
Condensates with different properties have significantly different growth patterns and coarsening kinetics. (A) A schematic for the use of a chemical dimerizer to induce condensate formation. (B) The ratio of the disordered protein condensate partition coefficient, viscosity, and surface tension to the coiled-coil condensate. (C,D) FRAP images and curves for a coiled-coil protein condensate (in C) and a disordered protein condensate (in D). Black lines are the exponential fits. Inset schematics are the predicted structure of the coiled-coil Mad1 protein (in C) and the disordered RGG domain of LAF-1 protein (in D) from AlphaFold2. (E,F) Fusion images and plots of aspect ratio over time for coiled-coil condensates (in E) and discorded protein condensates (in F). Black lines are the exponential fits. (G) Representative U2OS cell nucleus (magenta with DNA staining) containing condensates (green) formed by the coiled-coil protein imaged over time. Box indicates condensate that shrinks. (H) Condensate radius vs time for the six coiled-coil protein condensates shown in G. (I) Representative U2OS cell nucleus (magenta with DNA staining) containing disordered protein condensates (green) imaged over time. Boxes indicate condensates that shrink over time. (J) Condensate radius vs time for the six disordered condensates shown in I. (K) Fraction of growth types of the coiled-coil and the disordered protein condensates. Fusion events were scored as the coalescence of condensates, ripening events are characterized as the number of condensates shrinking while the remaining condensates grow, and diffusion-based growth is scored as continuous growth in the absence of ripening and can occur alongside fusion events (n.s., no significance; ***, p<0.001). (L) Change of average condensate radii over time. Condensate radius was normalized to the average condensate size at nucleation and time is normalized to the time nucleation occurs in the cell. Dashed lines are linear fits yielding indicated slopes. Scale bar, 5 *μ*m.

Having successfully induced two types of nuclear condensates, we tested whether their physical properties were significantly different. First, we estimated the partition coefficients by calculating the ratio of the mean fluorescent intensity in the condensed phase to the dilute phase. We found the partition coefficient of the disordered protein condensate to be half of that of the coiled-coil protein condensate, implying that the coiled-coil domains make for condensates with greater internal interaction strength than the disordered condensates (27) (Fig. 1B).

Next, we performed fluorescence recovery after photobleaching (FRAP) at the center of the condensates to compare the difference in viscosities, *η*, between the two condensates (22) (Fig. 1C,D). Exponential fits to the intensity recovery curves yield the recovery time *τ*, which for the coiled-coil condensate is in the order of minutes and that for the disordered condensate is in the order of seconds, suggesting that the two types of condensates exhibit significantly different dynamics (28) (Fig. 1C,D inserts). By combining *D* ∼ *r*^2^*/τ*, where *r* is bleached spot radius, with the Stokes-Einstein relation *D* = *k*_*B*_*T/*6*πηa*, where *k*_*B*_ is the Boltzmann constant, *T* is temperature, *η* is the viscosity, and *a* is the hydrodynamic radius of the diffusing particles that were assumed to scale linearly with the molecular weights of the phase separation proteins, we estimated that the coiled-coil condensate has a viscosity, *η*, 33 times larger than that of the disordered condensate (Fig. 1B).

We then used droplet fusion assays (4, 22, 28) to estimate the difference in surface tension, *γ*, of these two condensates. Similar to FRAP recovery, coiled-coil condensates take significantly longer time to round up during fusion (Fig. 1E,F). By assuming *η/γ* scales linearly with *τ/r* (4), where *r* is the length of the fusing condensates, and *τ* is the relaxation time obtained by using exponential fit of the relaxation curves, we estimated the surface tension, *γ*, for the disordered condensates to be 130 times greater than the coil-coil condensates (Fig. 1B).

Taken together, these analyses show that condensates formed with coiled-coil and disordered proteins have distinct material properties including viscosity and surface tension.

### Coiled-coil condensates and disordered condensates exhibit significantly different growth dynamics

Having confirmed the distinct properties of these two types of condensates, we proceeded to follow condensate coarsening dynamics. By using confocal microscopy to follow the condensates over time (Fig. 1G, I, Movie 1, 2) and by plotting the size of each condensate over time (Fig. 1H, J), we observed three modes of condensate coarsening: fusion, ripening, and continuous diffusion based growth, for both the coiled-coil and the disordered condensates (Fig. 1H,J).

By defining fusion as when two condensates coalesce to form a larger one, ripening as the shrinkage of condensates, and diffusion-based growth as continuous growth in the absence of ripening, we quantified the fraction of the different growth events by condensate type and observed significantly different growth patterns for the two condensates (Fig. 1K). First, ripening accounted for the majority of the growth events in disordered condensates compared to the least fraction in the coiled-coil condensates (Fig. 1K). In addition, the ripening time, defined as the time taken for a condensate to shrink over its radius, was 200 min/*μ*m for the disordered condensates and 1100 min/*μ*m for the coiled-coil condensates (Fig. S1). This suggests not only a greater ripening propensity for the disordered condensates but also faster ripening rates. Second, diffusion-based growth accounts for majority of growth for the coiled-coil condensates (67% of growth events) but it is only a small fraction for the disordered condensates (32% of growth events) (Fig. 1K). Lastly, growth by fusion for the disordered protein (20% of growth events) was similar to that of the coiled-coil condensate (27% of growth events) (Fig. 1K). Moreover, these growth patterns are also different from that reported for the FUS protein condensates where condensates grow dominantly by fusion while ripening and longer-term diffusion-based growth are absent (29, 30). These results suggest that condensate properties can affect the condensate growth patterns.

To assess the effect of condensate properties on coarsening rates, we plotted the average normalized condensate radius ⟨*R/R*_0_⟩ over time *t* (Fig. 1L). A power law fit following ⟨*R*⟩ ∼*t*^*β*^, where *β* is the growth exponent, yielded *β* = 0.22 for the coiled-coil condensates and *β* = 0.14 for the disordered condensate (Fig. 1L). Both exponents are smaller than predicted by theory (1*/*2 for diffusion based growth and 1*/*3 for fusion and ripening based growth) and are closer, even though still different from, the *β* = 0.12 observed for the FUS protein condensates (29). Given that the coiled-coil condensates and the disordered condensates had similar opportunities to grow by fusion but had different growth exponents (Fig. 1K, L), we suspect that their growth suppression may be attributed to the effect of chromatin on ripening or diffusion-based growth, different from the suppression of condensate fusion by chromatin for FUS condensates (29).

Together these results show that condensate growth is generally suppressed in the nucleus, but different condensates could be affected differently, likely due to different interplay between the condensates properties and the chromatin mechanics.

### Theoretical model of condensate growth in heterogeneous elastic media

To quantitatively understand how the different growth patterns emerge from the interplay between the condensate materials properties and the mechanics of the surrounding chromatin network, we developed a theory of condensate growth in elastic media (Fig. 2A-inset). Using the condition of material flux balance in and out of the condensate, we can formulate the growth dynamics of a spherical condensate in terms of its radius *R* as (31–33):

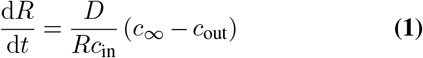

where *D* is the diffusion constant for the proteins in the dilute phase, *c*_∞_ is the far field concentration of the dilute phase, *c*_in_ and *c*_out_ are the concentrations of the proteins inside and outside the condensate. Considering the mass conservation of the total amount of condensate forming proteins and the effect of local pressure *P* on the dilute phase concentration, we arrive at (see Supplementary Materials Sec I for derivation):

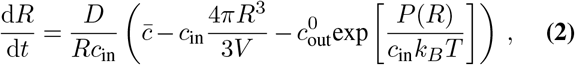

where 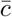 is the average protein concentration, 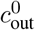 is the equilibrium protein concentration outside the droplet, *V* is the volume of the nucleus, and *T* is the temperature. Surface tension (*γ*) of the condensates and local stiffness of the chromatin network (*E*) both contribute to the condensate size-dependent pressure:

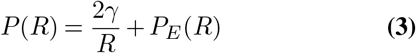

where *P*_*E*_ the pressure due to elastic deformation of the chromatin network. It has been suggested that chromatin networks exhibit hyper-elasticity due to their nonlinear stress-strain relationship (12, 34, 35). Consequently, we adopted a form for the mechanical pressure *P*_*E*_ derived from the known material response during the expansion of a cavity, created by condensate growth in a neo-Hookean elastic solid (36). This yields 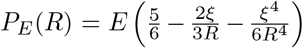, where *ξ* is the mesh size of the chromatin network surrounding the condensate.

**Fig. 2.**
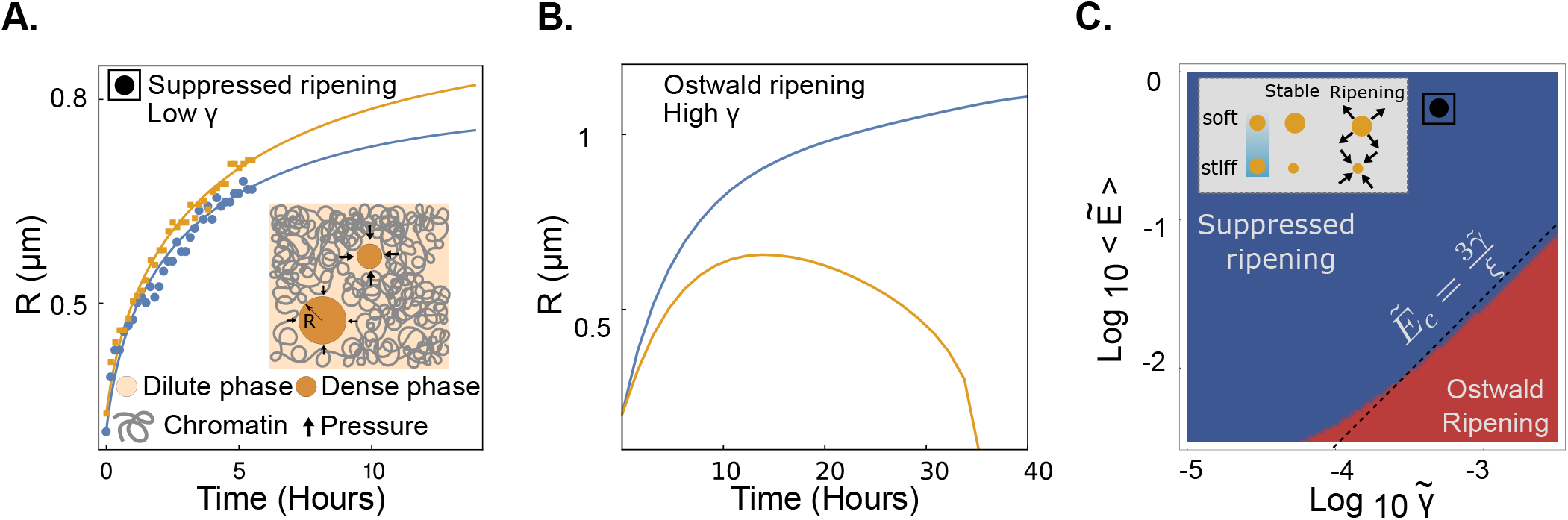
Distinct growth patterns emerge from the interplay between condensate surface tension and stiffness of the chromatin network. (A) Radii vs time for two condensates (labelled in yellow and blue) undergoing diffusive growth (suppressed ripening). The solid lines indicate model fits (see Methods for details) to the experimental data (solid circles). (B) Radii vs time for two condensates (labelled in yellow and blue) undergoing Ostwald ripening. Here the condensates have a larger surface tension compared to (A), with all other parameters fixed. (C) Phase diagram for condensate growth behavior as a function of normalized mean stiffness 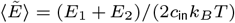 and surface tension 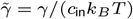, showing regimes of diffusive growth (stable) and Ostwald ripening. The parameter values used for panel A and B are 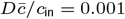, 4*πD/*3*V* = 10^−5^ *μ*m^−1^ s^−1^, 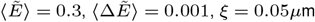 and the renormalized surface tension values are 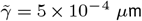 for panel A and 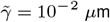 for panel B. For panel C the values are 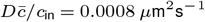 and 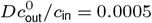 and other relevant parameters are same as in panel A & B.

The pressure *P*_*E*_ is zero when the condensate size is smaller than the mesh size (*R < ξ*).

Given the chromatin network is mechanically heterogeneous, its mesh size *ξ* and stiffness *E* are assumed to be local parameters (i.e., parameter value depends on the location of the condensate). On the other hand, condensate surface tension, diffusion constant, and the concentration values are assumed to be the same for all condensates within the nucleus. Properties of the condensate forming proteins and their molecular interactions determine the values of the parameters *γ, D, c*_in_, 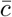 and 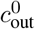. For *N* condensates growing in a shared environment, the growth dynamics for the *i*^*th*^ condensate can be written as:

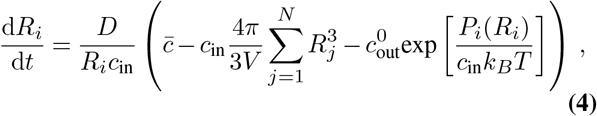

where *R*_*i*_ is the size of the *i*^*th*^ condensate, and *P*_*i*_ is the local pressure around the *i*^*th*^ condensate with stiffness *E*_*i*_ and mesh size *ξ*_*i*_.

### Model reveals the relative roles of surface tension and chromatin elasticity on condensate growth dynamics

We first consider the growth of two condensates of radii *R*_1_ and *R*_2_, embedded in the chromatin network with local stiffness values *E*_1_ and *E*_2_, respectively. This simplified two-droplet model is useful to elucidate the underlying mechanism of condensate ripening and its suppression, regulated by the interplay between droplet surface tension and chromatin stiffness. We used the growth model (Eq. 4) to fit experimental data from coiled-coil condensates showing suppressed ripening (Fig. 2A). Details of the fitting method is provided in the Methods section. Theoretical results show that the condensates can undergo Ostwald ripening when the surface tension is increased keeping all other parameters and the stiffness of the surrounding elastic medium the same (Fig. 2B). This explains the increased ripening of the disordered condensates (Fig. 1K) that have a higher surface tension compared to the coiled-coil condensates (Fig. 1B), with the latter exhibiting mostly diffusive growth (Fig. 1K). The theoretically obtained phase diagram in the plane of renormalized average stiffness 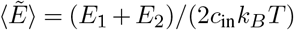) and surface tension 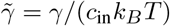, with a fixed stiffness difference 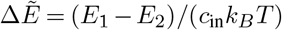, shows the parameter regimes of suppressed ripening and Ostwald ripening phases, indicating that protein property 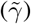 and the mechanical property of the chromatin network 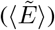 can determine the resulting condensate growth dynamics (Fig. 2C).

### Condensate material properties and chromatin heterogeneity determine the modes of ripening

The low surface tension in coiled-coil condensates precludes Ostwald ripening as the mechanical pressure from the surrounding elastic network will stabilize the ripening droplets at different sizes (Fig S1, Fig. 2A). However, we observed a small fraction of coiled-coil condensates undergoing ripening in the elastic chromatin network (Fig. 1G,H&K). Recent studies on oil droplets in silica gel have shown that droplets can undergo elastic ripening due to differences in mechanical pressure from the surrounding medium (37). However, to the best of our knowledge, no instances of elastic ripening have been reported in living cells. The fitting of our model to the experimental data of ripening in coiled-coil condensates indicates a significant difference in local stiffness values which might be enough to drive elastic ripening of the coiled-coil condensates (Fig S2).

To investigate the underlying mechanism of condensate ripening in the chromatin network with spatially heterogeneous mechanical properties, we derived a linearized theory for the dynamics of the droplet size difference ∆*R* given by:

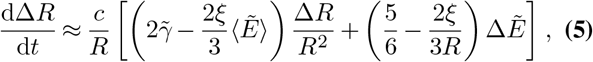

where *R* is the typical condensate size (see Supplementary Materials Sec II for details). The linearized theory is valid when ∆*R* is small, such that *R*_1_ ∼*R*_2_ ∼*R*, and is useful to derive the condition for the onset of ripening. Stability analysis of Eq. 5 reveals three distinct modes of condensate coarsening. For large enough surface tension 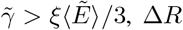 increases over time, leading to the growth of the larger condensate at the expense of the smaller one, suggestive of Ostwald ripening (Fig. 3A). This is comparable to the disordered condensates in our experiments that possess high surface tension and exhibit Ostwald ripening (Fig. 1K,L). By contrast, when 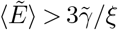, Ostwald ripening is suppressed by the mechanical pressure from the surrounding chromatin network, leading to a stable size difference between the condensates 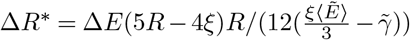 (see Fig. 3A).

**Fig. 3.**
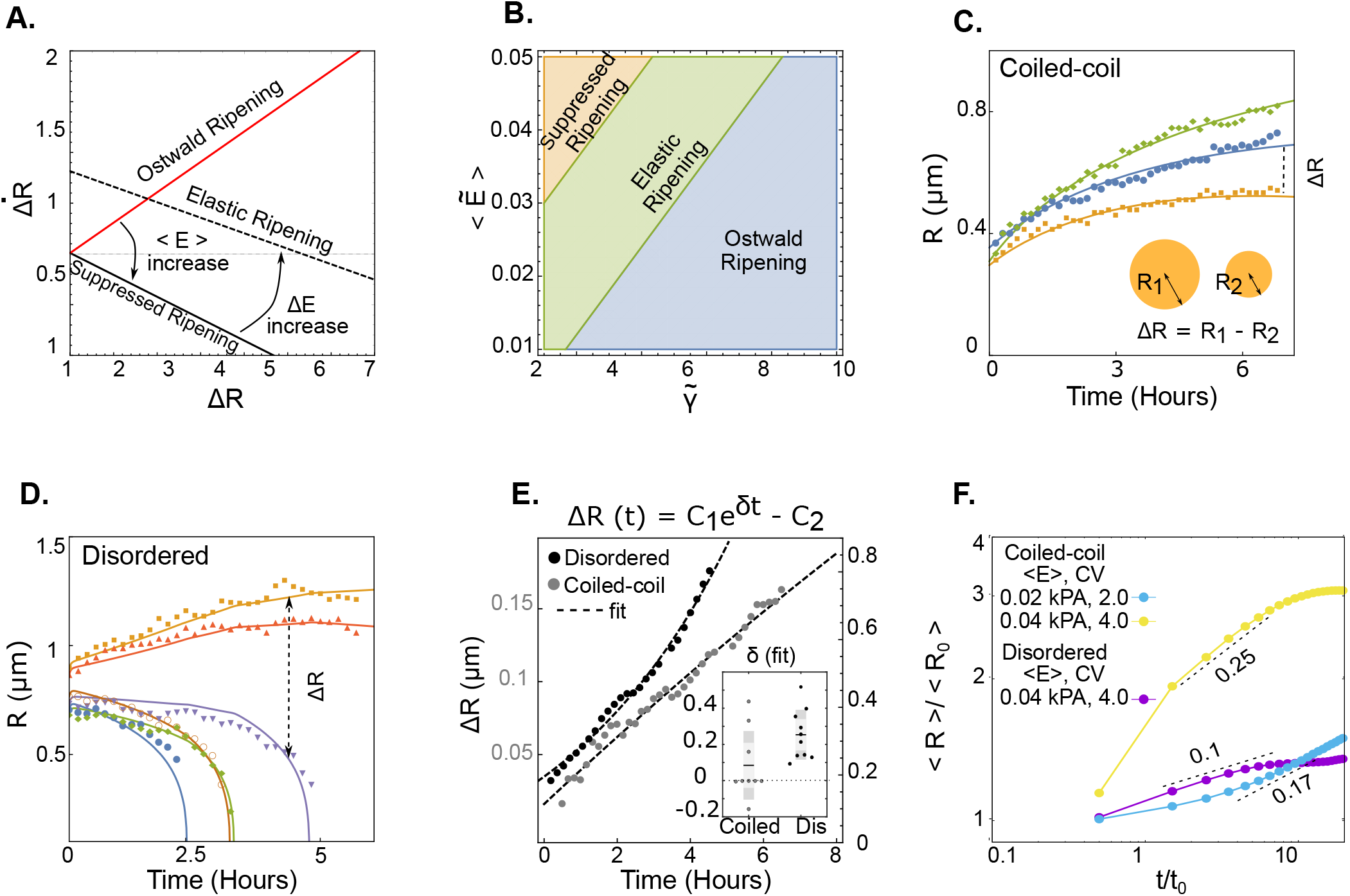
Mechanical heterogeneity of the surrounding elastic network induces elastic ripening and slow growth of condensates. (A) Phase portrait showing the dependence of time-derivative of the condensate size difference, 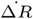, as a function of ∆*R*. Three distinct growth patterns emerge depending on the slope and the intercept of 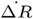 vs ∆*R*. (B) Phase diagram in the plane of surface tension 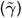 and mean stiffness 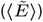 showing the parameter regimes for suppressed ripening, Ostwald ripening and elastic ripening. (C) Time evolution of the sizes of three growing coiled-coil condensates, exhibiting suppressed ripening. Solid lines are model fits to the experimental data. (D) Time evolution of the sizes of five disordered condensates, showing instances of ripening. Solid lines are model fits to the experimental data. (E) ∆*R* vs time for ripening droplets in disordered and coiled-coil condensates show qualitatively distinct trends. (Inset) Characterization of ripening events by obtaining the rate *δ* (in units of hour^−1^) by fitting linear theory (Eq. 5) to experimental data. (F) The numerical solution of the model (Eq. 4) for multiple condensates growing in a heterogeneous stiffness landscape predicts power-law scaling of mean condensate sizes during growth. We use high and low values of mean stiffness (⟨*E*⟩) and coefficient of variation (*CV*_*E*_) for proteins with low (coiled-coil) and high (disordered) surface tension to predict how different stiffness distributions affect condensate growth. Here ⟨*R*_0_⟩ is the mean initial radius of the condensates and the characteristic timescale *t*_0_ = 600 seconds. The parameters used are provided in Table. 1.

In a homogeneous elastic medium (i.e., ∆*E* = 0), the suppression of ripening will result in equal sized condensates (∆*R*^∗^ = 0) as has been previously observed (37). Inhomogeneity in the elastic environment will give rise to stable condensates of different sizes (bigger condensates at regions of lower stiffness) and indeed we observe this in the case of suppressed ripening (Fig. 1H). Interestingly, This difference in size between the stable condensates (∆*R*^∗^) increases with increasing stiffness difference ∆*E* and can lead to complete loss of one of the condensates in the stiffer environment when ∆*R*^∗^ ≥ *R*. We identify this ripening event as elastic ripening driven by the mechanical pressure difference between the condensates due to a difference in their local stiffness values (Fig. 3A). In the case of coiled-coil condensates that have very low surface tension, we do observe a few cases of ripening where Ostwald ripening is unlikely (Fig. 1G,H). We used Eq. 5 to predict the phase diagram of the ripening behaviors (at constant 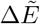) as a function of 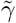 and 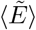 (Fig. 3B). The phase diagram shows a transition from Ostwald ripening to elastic ripening and suppressed ripening as surface tension is reduced and stiffness is increased.

We fit our model (Eq. 4) to experimental data to gain quantitative insights into the kinetics of condensate growth and ripening (Fig. 2A, Fig. 3C-D). In particular, the model can also be utilized to infer the mode of ripening (Ostwald vs elastic ripening) directly from the experimental data. The linearized theory predicts that the dynamics of the condensate size difference ∆*R* evolves in time as

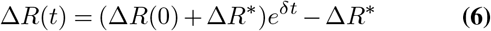

where 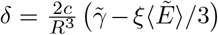. For Ostwald ripening *δ >* 0 and for elastic ripening (as well as for diffusive growth) *δ <* 0 (Fig. 3A). The quantity ∆*R* can be easily extracted from the experimental data (Fig. 3C,D) and we can fit the time evolution of ∆*R* to find *δ* for both coiled-coil and disordered condensates (Fig. 3E). Our analysis shows *δ <* 0 for the majority of the ripening cases in coiled-coil condensates indicating elastic ripening. In contrast, we find that *δ >* 0 for all of the ripening cases in disordered condensates, indicating Ostwald ripening (Fig. 3E-inset). While the theoretical prediction combined with the trend in ∆*R* data from the experiments indicates elastic ripening in coiled-coil condensates, the magnitudes of the ripening rate *δ* are small (Fig. 3E-inset) such that complete dissolution of coiled-coil condensates is not observed over the experimental timescale (Fig. 3C). The slow rate of elastic ripening may result from the smaller values of 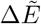, as the coiled coil condensates most likely nucleate in regions of low stiffness in the chromatin network.

### Effect of mechanical heterogeneity on condensate nucleation and growth

The chromatin network inside the nucleus is spatially heterogeneous and the local mechanical properties are determined by the local density and the architecture of the chromatin network (38, 39). To understand how heterogeneity in chromatin elasticity affects condensate nucleation and growth, we extended our theory to incorporate the effects of medium elasticity on condensate nucleation. The probability *p*_nuc_ of nucleating a droplet of size *R*_0_ is given by *p*_nuc_ ∝ exp(− ∆*G/k*_*B*_*T*), where ∆*G* is the free energy change due to nucleation:

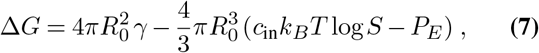

with *S* the extent of super-saturation (see Supplemental Materials Sec. III and Fig. S3). Upon nucleation, the growth dynamics of multiple condensates are given by the system of equations in Eq. 4. The chromatin stiffness landscape is determined by sampling the local stiffness from a normal distribution with mean 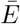 and coefficient of variation *CV*_*E*_. We considered a finite pool of condensate material such that the level of supersaturation *S* decreases as more condensates are nucleated. Condensate nucleation, growth and coarsening were then studied for different stiffness distributions, for both the coiled-coil and disordered condensates.

In the absence of a surrounding elastic medium, the average condensate size grows in time following the scaling law ⟨*R*⟩∼ *t*^1*/*3^, as expected for coarsening via Ostwald ripening (31, 32). In the presence of an elastic medium, ⟨*R*⟩ does not adhere strictly to a power law scaling over time, but one can fit a power law during the growth phase (Fig. 3F). We quantified the scaling behavior of condensate growth with time as, ⟨*R*⟩ */* ⟨*R*_0_⟩ ∼*t*^*β*^, where *R*_0_ is the initial condensate size and *β* is the scaling exponent. In particular, we computed how *β* depends on the mean and the variance in stiffness of the surrounding elastic medium, for both coiled-coil and disordered condensates (Fig S4).

We found that the coiled-coil condensates grow faster than the disordered condensates, with *β* = 0.25 for coiled-coil and *β* = 0.1 for disordered condensates (Fig. 3F), in reasonable quantitative agreement with the experimental data (Fig. 1L). Further theoretical analysis predicts that the scaling exponent *β* decreased with decreasing *CV*_*E*_ and increased with increasing 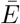 for low surface tension condensates (Fig. 3F, Fig S4), while *β* did not change significantly with stiffness variations for high surface tension condensates (Fig S4). This result contrasts with the known scaling behaviors of condensate coarsening in liquids where the scaling exponent is independent of the properties of the liquid.

### Chromatin heterogeneity promotes the growth of low surface tension condensates

To test the model prediction on the effects of chromatin mechanical heterogeneity on the growth of different condensates, we assessed condensate growth in cells with different chromatin environments. First we treated U2OS cells with Trichostatin-A (TSA), a histone deacetylase inhibitor that has been shown to de-condense chromatin and soften the nucleus (40). In addition, we switched cell type to HeLa, which has different nuclei size and chromatin organization. The difference in the chromatin organization in the three types of cells is estimated by differences in the chromatin intensity distribution (Fig. 4A,B). HeLa nuclei have significantly greater mean chromatin intensity and variance than untreated U2OS cells, while TSA treatment lowered the chromatin mean intensity and variance in U2OS cells as expected from the de-condensation of chromatin and the resulting increase in the homogeneity of the chromatin environment (Fig. 4C, S5A). This suggests both the mean chromatin stiffness and the variance could be different in these three types of cells, reflecting different chromatin environments.

**Fig. 4.**
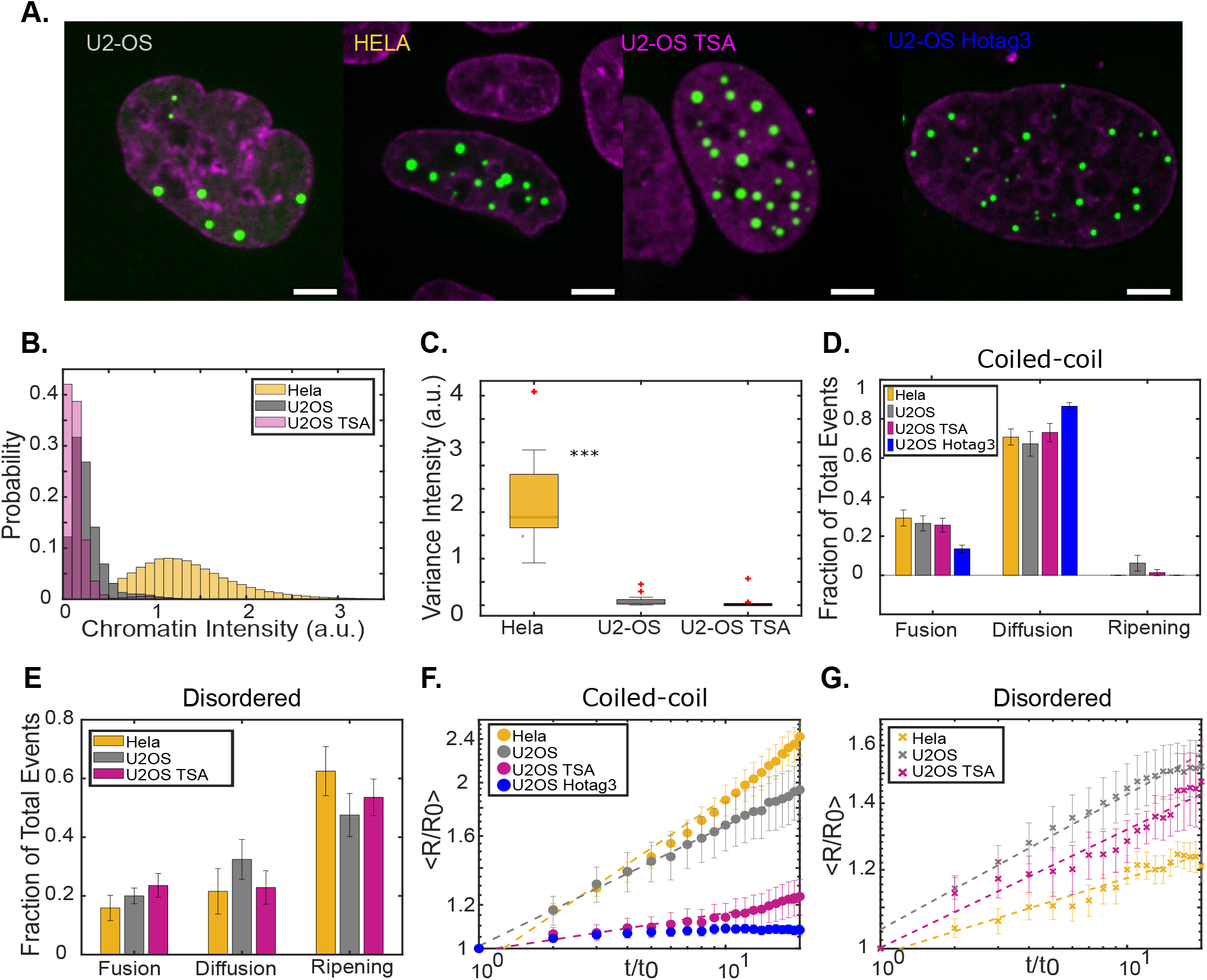
Chromatin heterogeneity affects the growth of coiled-coil condensates more than disordered condensates. (A) Images of representative nuclei (magenta with SPY650DNA staining) with coiled-coil condensates (green) nucleated with LacI in a U2OS cell, a HeLa cell, and a TSA-treated U2OS cell, and that nucleated with Hotag3 in a U2OS cell (scale bar, 5 *μ*m). (B) Distributions of the chromatin intensity for HeLa, U2OS, and TSA treated U2OS cells. (C) The variance of chromatin intensity for HeLa, U2OS, and, TSA treated U2OS cells. (D) Quantification of the growth types of coiled-coil condensates nucleated with LacI in HeLa, U2OS, and TSA treated U2OS cells, and that nucleated with Hotag3 in U2OS cells. (E) Quantification of the growth types of the disordered protein condensates in HeLa, U2OS, and TSA treated U2OS cells. (F) The average radii over time for coiled-coil condensates nucleated with LacI in HeLa, U2OS, TSA treated U2OS cells, and that nucleated with Hotag3 in U2OS cells. (G) The average radii over time for disordered condensates in HeLa, U2OS, TSA treated U2OS cells. In (F) and (G), the radii were normalized to the initial average droplet size by the cell and the time was normalized to the time condensates were nucleated.

Despite the significant difference in chromatin organization in these nuclei, the growth patterns of the coiled-coil and the disordered condensates remained similar (Fig S5B,E). The coiled-coil condensates still grew mainly by diffusion (73% in TSA treated U2OS cells and 71% in HeLa cells) while the disordered protein condensates grew mostly through ripening (54% in TSA treated U2OS cells, 63% in HeLa cells) (Fig. 4D,E). This is consistent with our theoretical results that changes in mean chromatin stiffness or the variance did not impact the pattern of coarsening for coiled-coil and disordered condensates (Fig. S4B). The difference in chromatin environment did not have a significant effect on the growth by fusion for both the condensate types, suggesting that the condensate mobility was similarly limited by chromatin if at all (Fig. 4D,E). For the coiled-coil condensates, the reduction of ripening in TSA treated U2OS cells (1.3%) and in HeLa cells (none) is not conclusive due to the scarcity of ripening events (Fig. 4D). Comparisons of the ripening time of disordered condensates in the different chromatin environments showed that the chromatin environment did not have a significant impact on ripening dynamics (Fig S5F). These data suggest that condensate growth patterns are dominated by condensate properties and are not significantly affected by physical changes in the chromatin environment.

However, differences were observed in the condensate growth rates (Fig. 4F,G). For coiled-coil condensates, the growth exponent *β* decreased from 0.22 to 0.07 in TSA treated U2OS cells and increased to 0.3 in HeLa cells, which is still lower than 0.5 predicted by diffusion based growth (29). For disordered condensates, the growth exponent *β* reduced slightly from 0.14 in untreated cells to 0.12 in TSA treated U2OS cells and 0.08 in HeLa cells (Fig. 4G). These trends are consistent with theoretical predictions (Fig S4). Overall, more changes are observed for the coiled-coil condensates and the changes in response to TSA treatment and cell type is different for the two condensates, suggesting that different condensates are affected differently by changes in chromatin landscape.

In addition to the marked difference in the growth exponents, we observed that, after TSA treatment, the average number of condensates formed by the coiled-coil protein significantly increased from 8 condensates per cell to 18 condensates, while the count for the disordered condensates remained similar (8 condensates per cell) (Fig S5G). We hypothesized that TSA treatment may favor coiled-coil condensate nucleation, forming more condensates whose competition for growth quickly quenched available material, resulting in the suppressed growth. To test this hypothesis, we used the stronger nucleator, hexamer Hotag3, that was used for the disordered proteins to induce coiled-coil condensates in untreated U2OS cells (Fig. 4A). As expected, more condensates were formed than with the LacI nucleator and the condensate count (19 condensates per cell) was comparable to that in TSA treated cells (Fig S5G), agreeing with the previous findings that condensate nucleation in cells can be modulated by molecular interactions and cellular processes (41). Similar to LacI-induced coiled-coil condensates in various chromatin environments, Hotag3-induced coiled-coil condensates still mainly grow by diffusion (Fig. 4D), suggesting that the growth pattern is not sensitive to nucleation. However, these condensates have the smallest growth exponent of 0.02, closer to that in the TSA treated cells (Fig. 4F). This suggests that chromatin heterogeneity can control growth dynamics of the low surface tension condensates by affecting nucleation.

## Discussion

Nuclear condensates have been previously shown to coarsen primarily via coalescence (29). Here we report that nuclear condensates can coarsen via coalescence, diffusive growth, Ostwald or elastic ripening, and the prevalence of each mode is dictated by the material properties of the condensate. Similar to previously reported suppression of growth in elastic environment (18, 19, 29), we also observed reduced coarsening rates for condensates with different materials. Theoretical modeling shows that different patterns of condensate growth and their coarsening rates result from the interplay between condensate surface tension and chromatin stiffness. This indicates that cells can modulate both condensate properties and chromatin stiffness to regulate condensate growth in the nucleus. After DNA damage, for example, it is known that chromatin is softened and DNA damage proteins undergo multiple types of post-translational modifications (42). It is possible that some of these changes are to regulate chromatin stiffness and condensate properties so that condensate growth at DNA damage sites is favored over other chromatin regions for efficient DNA damage repair.

Many membrane-less compartments need to be stabilized after nucleation. For phase separation to be the driver of the formation of these compartments, an outstanding question has been how do cells prevent Ostwald ripening to achieve compartment stabilization (43). Previous work has shown that internal biochemistry can help prevent Ostwald ripening, which was used to explain centrosome stabilization (8). In addition, for P granules, protein clusters absorbed on the surface can prevent their Ostwald ripening via Pickering stabilization (44). In the human cell nucleus, Ostwald ripening of FUS protein condensates was inhibited by the elastic chromatin (45). By comparing ripening of the two types of condensates in human cell nucleus, our work shows that the elastic chromatin network can stabilize condensates against Ostwald ripening but only when condensate surface tension is low. This suggests that protein interactions might have been evolved to generate low surface tension condensates to counteract Ostwald ripening. Indeed, there are a wide range of interaction types/strengths from various types of domains/motifs that biological molecules can explore to form condensates (46). Much effort has been focused on how these interactions drive the phase separation process. Our work suggests that understanding how these molecular interactions lead to condensates with different material properties are also important for understanding condensate growth and size control in the nucleus.

By observing condensate growth in different chromatin environments, we discovered that patterns of coarsening show little dependence on the changes in chromatin properties. However, we observed that condensate growth rates are significantly different for different condensates, agreeing with our model prediction that low surface tension condensates grow faster in heterogeneous environments than high surface tension condensates. Interestingly, we also observed noticeable differences in nucleation in different chromatin environment that contributes to the difference in growth dynamics. Previous work has shown that nucleation affects condensate size distribution in cells (47). Our findings imply that nucleation landscape also affects condensate growth kinetics in the nucleus, highlighting the importance of nucleation control for biomolecular condensates. Much like growth, the material properties of condensates could also be significant factors influencing the nucleation landscape within the nucleus. It is possible that cells have adapted to utilize the interaction between chromatin mechanics and condensate properties to regulate nucleation for various biological functions.

## Methods

### Plasmids

The plasmids used to make the disordered protein condensates were NLS-3xHalo-GFP-Hotag3 and RGG-mCherry-RGG-eDHFR (23, 25, 26). The plasmids used to make the coiled-coil protein condensates were mScarlet-eDHFR-Mad1 as the phase separating protein (26) and Halo-GFP-LacI-NLS as the anchor protein.

### Cell culture

The U2OS cells were gifted by Dr. Eros Lazzerini Denchi and the HeLa cells used were recombination-mediated cassette exchange (RMCE) acceptor cells (50). The HeLa cells were cultured in growth medium (Dulbecco’s Modified Eagle’s medium with 10% FBS and 1% peni-cillin–streptomycin) and the U2OS cells were cultured in the same media supplemented with 1% glutamine. All cells were cultured at 37 ºC in a humidified atmosphere with 5% CO2. Cells were seeded on 22×22 mm glass coverslips (no. 1.5; Fisher Scientific) coated with poly-D-lysine (Sigma-Aldrich) for transfection at 60 − 70% confluence using Lipofectamine 2000 (ThermoFisher Scientific) according to protocols detailed in published works (23, 25). For the disordered condensate experiments, the cells were co-transfected with 0.5 μg of the NLS-3xHalo-GFP-HoTag3 plasmid DNA and 1 μg of the RGG-mCherry-RGG-eDHFR plasmid. The coiled-coil condensate experiment co-transfections were 0.5 μg of the Halo-GFP-LacI-NLS and 1 μg mScarlet-eDHFR-Mad1. The phase-separating protein and anchor protein combination constructs were thus transiently expressed for 24-48 hours prior to imaging. In the instance of chromatin perturbation, transfected cells (24 hours) were treated with 0.2 mg/ml of Trichostatin-A (TSA, Sigma-Aldrich) for 24 hours before imaging.

### Image acquisition and dimerization

For live imaging, the coverslips were mounted into magnetic chambers with 1ml of growth media and 1x SPY650DNA (Spirochrome) as a nuclear stain. Cells without condensates, but with high expression of both the anchor and the phase separating protein were selected and image acquisition began. At the end of the first time loop, an additional 500 μl of growth media containing the chemical dimerizer TMP-Fluorobenzamide-Halo (TFH) to a final condentration of 100 μM was added to the stage to induce condensate formation in the mounted cells expressing RGG- mCherry-RGG-eDHFR and NLS-3xHalo-GFP-HoTag3 or mScarlet-eDHFR-Mad1 and Halo-GFP-LacI-NLS (23, 25). Z-stack images were collected with 0.6 μm spacing for a total depth of up to 12 μm, at 10 minute intervals for 6-12 hours. A Nikon Eclipse Ti2 microscope with a Tokai Hit stage incubator (with CO2), a Yokogawa CSU-X1 spinning disk confocal equipped with a 100x/1.4 NA oil immersion objective, a XY Piezo-Z stage (Applied Scientific Instrumentation), 488 nm (GFP), 561 nm (mScarlet), and 647 nm (Cy5) laser modules and an EMCCD camera was used to obtain the time lapse images (23, 25).

### Image processing

Maximum projections of the z-stack images were pre-processed and the condensates segmented using NIS-Elements Advanced Research Analysis software (5.30.05 64bit, Nikon). The nuclear stain channel (Cy5) was used to isolate nuclear binaries which were tracked and measured using the NIS Elements tracking module. The nuclear binary tracks were then exported to excel spread sheets which were compiled using MATLAB (R2020b, Mathworks) for growth dynamics plots.

**Table 1.**
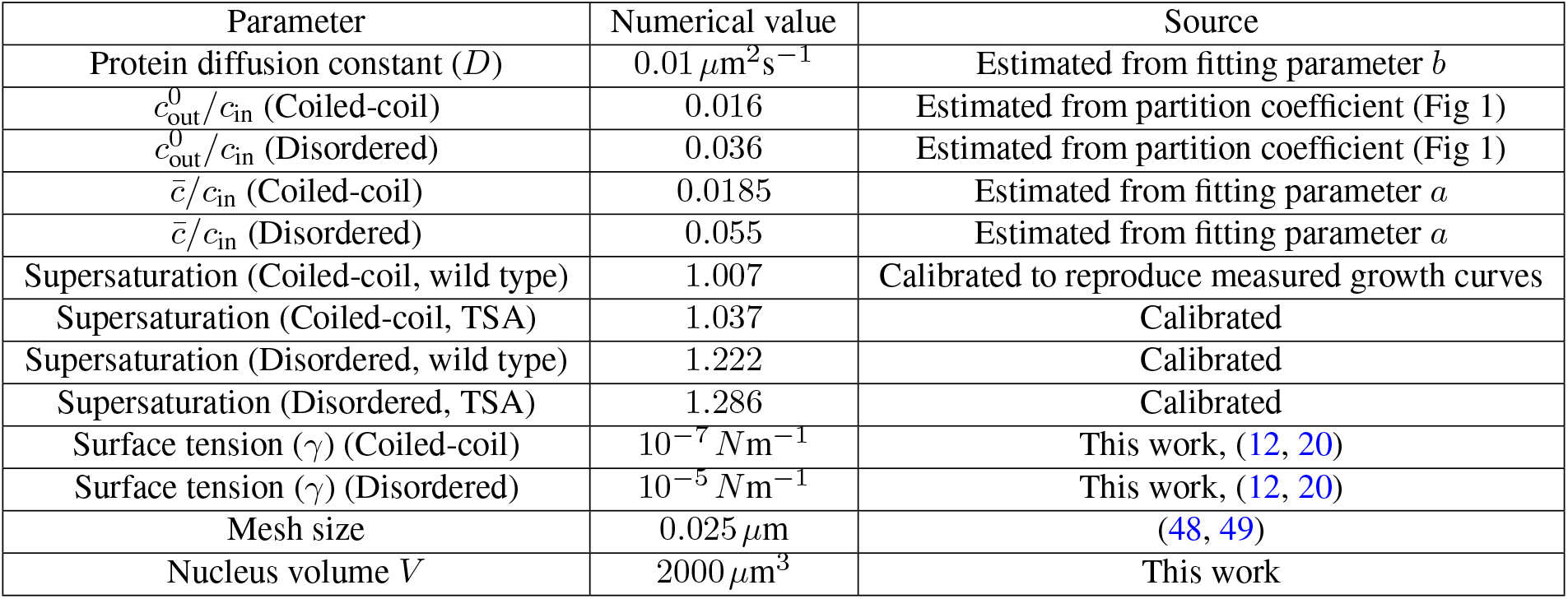
Parameter values used in numerically solving the model.

| Parameter | Numerical value | Source |
| --- | --- | --- |
| Protein diffusion constant ($D$) | $0.01 \mu\text{m}^2\text{s}^{-1}$ | Estimated from fitting parameter $b$ |
| $c_{\text{out}}^0/c_{\text{in}}$ (Coiled-coil) | 0.016 | Estimated from partition coefficient (Fig 1) |
| $c_{\text{out}}^0/c_{\text{in}}$ (Disordered) | 0.036 | Estimated from partition coefficient (Fig 1) |
| $\bar{c}/c_{\text{in}}$ (Coiled-coil) | 0.0185 | Estimated from fitting parameter $a$ |
| $\bar{c}/c_{\text{in}}$ (Disordered) | 0.055 | Estimated from fitting parameter $a$ |
| Supersaturation (Coiled-coil, wild type) | 1.007 | Calibrated to reproduce measured growth curves |
| Supersaturation (Coiled-coil, TSA) | 1.037 | Calibrated |
| Supersaturation (Disordered, wild type) | 1.222 | Calibrated |
| Supersaturation (Disordered, TSA) | 1.286 | Calibrated |
| Surface tension ($\gamma$) (Coiled-coil) | $10^{-7} \text{Nm}^{-1}$ | This work, (12, 20) |
| Surface tension ($\gamma$) (Disordered) | $10^{-5} \text{Nm}^{-1}$ | This work, (12, 20) |
| Mesh size | $0.025 \mu\text{m}$ | (48, 49) |
| Nucleus volume $V$ | $2000 \mu\text{m}^3$ | This work |

### Model Fitting

The dynamic equation for droplet growth was reparametrized to fit experimental growth curves. We rewrite Eq. 4 as:

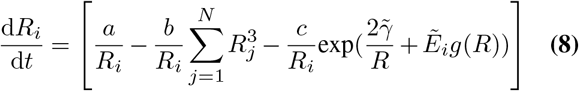

where the renormalized surface tension and stiffness are defined as 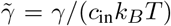 and 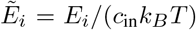 respectively. The other parameters are 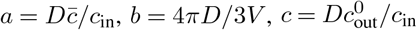, and 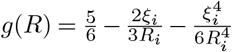. At room temperature *k*_*B*_*T* ∼4.114 ×10^−21^ *J* and *c*_in_ ∼10^5^ *μm*^−3^ (51, 52) leading to *c*_in_*k*_*B*_*T* ∼4 ×10^−10^ *N μm*^−2^. Depending on the condensate forming protein, the surface tension may be in a range ∼10^−7^ −10^−4^ *Nm*^−1^ (12, 20) and using the value of *c*_in_*k*_*B*_*T* ∼4 ×10^−10^ *N μm*^−2^ we can estimate the range of the parameter 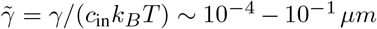. Using the same value for *c*_in_*k*_*B*_*T* in a chromatin stiffness range of 1 −1000 Pa (39, 53), the parameter 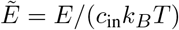 will have values in the range ∼0.001 −1.

We estimated the diffusion constant for the dilute phase to be *D* = 0.01 *μm*^2^*sec*^−1^ and obtained the value of parameter *c* as 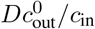 by using the ratio of protein intensities outside and inside the condensate to estimate 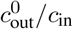. We found the mean value of 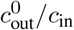 to be 0.016 and 0.036 for coiled-coil and disordered proteins, respectively. The mesh size was considered to be *ξ* = 0.025 *μm* (48, 49) and the surface tension 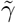 was taken to be 0.0001 and 0.01 for the coiled-coil and disordered protein condensates, respectively.

We used a global optimization algorithm *multistart* combined with a nonlinear curve fitting method *lsqcurvefit* in MAT-LAB (54) to find the parameters *a, b* and the local stiffness values 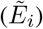 for each droplet growing in a cell. The algorithm used multiple start points to sample multiple basins of attraction and solve a local optimization to find a set of parameters ({*a, b, E*_*i*_} corresponding to the local start point) that minimized the error 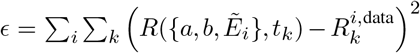, where *R*^*i*,data^(*t*) is the experimentally measured size of the *i*^*th*^ condensate at time *t* and *R*(*t*) is the theoretical prediction of *i*^*th*^ condensate size obtained from solving Eq. 8 for all condensates in a cell. The sum with index *i* and *k* indicate summation over all condensates in a cell and all time points of observation respectively. We used 200− 500 iterations of multi-start to find a convergent trend in the error and considered the fitting parameters corresponding to the smallest *ϵ*.

## Supplementary Movies

**Movie S1. Growth of coiled-coil condensates**. Condensates were formed with Halo-GFP-LacI-NLS (green) and mScarlet-eDHFR-Mad1 (not shown) by adding dimerizer TFH at the end of the first time loop. U2OS nuclei (magenta) were stained with SPY650DNA. For Fig. 1G,H.

**Movie S2. Growth of disordered condensates**. Condensates were formed with NLS-3xHalo-GFP-Hotag3 (green) and RGG-mCherry-RGG-eDHFR (not shown) by adding dimerizer TFH at the end of the first time loop. U2OS nuclei (magenta) were stained with SPY650DNA. For Fig. 1I,J.

## Acknowledgements

SB and DC acknowledge support from the National Institutes of Health (NIH R35 GM143042 to SB, GM118510 to DC) and HZ acknowledges support from the National Science Foundation (NSF MCB-2145083). SB, HZ, DB and TC gratefully acknowledge support from the David Scaife Foundation. JK and TC acknowledge support from NIH T32 Fellowship (T32GM133353). TC also acknowledges support from the Brian and Diane Smith Fellowship, GEM Fellowship, CIT Presidential Fellowship, Mahmood I. Bhutta Fellowship.

