## Supplemental Text and Figures for "Interplay of condensate material properties and chromatin heterogeneity governs nuclear condensate ripening"

### Supplementary Materials for
### Interplay of condensate material properties and chromatin heterogeneity governs nuclear condensate ripening

Deb Sankar Banerjee,[1,*] Tafadzwa Chigumira,[2,*] Rachel M. Lackner,[3] Josiah Kratz,[4] David M. Chenoweth,[3] Shiladitya Banerjee,[1,†] and Huaiying Zhang[4,2,†]

[1]*Department of Physics, Carnegie Mellon University, Pittsburgh, PA 15213, USA*

[2]*Department of Chemical Engineering, Carnegie Mellon University, Pittsburgh, PA 15213, USA*

[3]*Department of Chemistry, University of Pennsylvania, Philadelphia, PA 19104, USA*

[4]*Department of Biological Sciences, Carnegie Mellon University, Pittsburgh, PA 15213, USA*

[*] equal contribution

### I. THEORY OF CONDENSATE GROWTH IN ELASTIC MEDIA

Here we detail the derivation of the dynamical equations governing condensate size and discuss how the presence of the elastic media affects condensate coarsening dynamics.

#### A. Growth of a single condensate

We first consider the growth of a single condensate in a homogeneous elastic network with stiffness $E$. We assume that the condensate-forming proteins can diffuse freely through the network mesh in the dilute phase, while the dense phase can deform the network locally as they grow to be larger than the average network mesh size $\xi$. The dynamics of the protein concentration field, $c$, is given by the diffusion equation:

$$\partial_t c = D\nabla^2 c \,, \tag{S.1}$$

where the protein diffusivity $D$ is taken to be the same for both the dilute phase outside the condensate and the dense phase inside the condensate. Assuming the system to be in a quasi-static state where diffusion is much faster compared to growth, we can solve for the steady-state concentration field using the equation

$$\nabla^2 c \simeq 0 \,, \tag{S.2}$$

with the boundary conditions given by Gibbs-Thompson relation and $\partial_r c \mid_{r=0} = 0$. We consider a perfectly spherical condensate, a fluid droplet of radius $R$. The solution inside the condensate is given by $c = c_{\text{in}}$. Whereas outside the condensate ($r > R$), the concentration field is given by:

$$c(r) = c_\infty + (c_{\text{out}} - c_\infty)\frac{R}{r} \,. \tag{S.3}$$

where $c_\infty$ is the far-field concentration, assumed to be constant. Growth of the droplet is dictated by the net influx of material into the condensate. A local material conservation leads to the equation:

$$c_{\text{in}}\Delta V_d = \int_{S_d} (j_{\text{in}} - j_{\text{out}})dA\,\Delta t \,, \tag{S.4}$$

where $j_{\text{in}}$ and $j_{\text{out}}$ are the normal components of the inward and outward diffusive fluxes given by $j_{\text{in/out}} = -D\nabla c_{\text{in/out}}$, and the integral is over the surface of the condensate $S_d$. Assuming homogeneous growth, we arrive at the equation:

$$c_{\text{in}}\,\Delta R = (j_{\text{in}} - j_{\text{out}})\,\Delta t \tag{S.5}$$

where we have taken $\Delta V = 4\pi R^2 \Delta R$. In the limit $\Delta t \to 0$ we get

$$\frac{dR}{dt} = \frac{j_{\text{in}} - j_{\text{out}}}{c_{\text{in}}} \,. \tag{S.6}$$

We can calculate the diffusive fluxes from the solution of $c(r)$ and get $j_{\text{in}} = 0$ and $j_{\text{out}} = \frac{D}{R}(c_{\text{out}} - c_\infty)$. The growth rate of the droplet can now be derived from Eq. S.6:

$$\frac{dR}{dt} = \frac{D}{Rc_{\text{in}}} \left(c_\infty - c_{\text{out}}\right) \ . \tag{S.7}$$

We assume that the growing droplets are in equilibrium with their surrounding, which exerts a pressure $P$ onto the droplet. This pressure has two contributions - (i) Laplace pressure $2\gamma/R$ due to surface tension $\gamma$ of the dense phase, and (ii) a mechanical pressure $P_E$ exerted by the surrounding network. We can estimate the effect of the exerted pressure $P$ on the equilibrium densities ($c_{\text{in}}$ and $c_{\text{out}}$) from a Maxwell construction. Assuming equilibrium and strong phase segregation ($c_{\text{in}} \gg c_{\text{out}}$), the chemical potential $\mu$ is related to the pressure as

$$P = c_{\text{in}}(\mu_{\text{out}} - \mu^0_{\text{out}}) \tag{S.8}$$

where $\mu^0_{\text{out}}$ is the chemical potential in the absence of pressure. Assuming an ideal dilute phase, the chemical potential is related to concentration as $\mu(c_{\text{out}}) = k_B T \ln(c_{\text{out}})$, where $k_B$ is the Boltzmann constant and $T$ is the absolute temperature. Using the above relations we can arrive at the expression of concentration outside the droplet surface given by

$$c_{\text{out}} = c^0_{\text{out}} \exp\left(\frac{P}{c_{\text{in}} k_B T}\right) \ . \tag{S.9}$$

The pressure $P$ is given by

$$P = \frac{2\gamma}{R} + E\left(\frac{5}{6} - \frac{2\xi}{3R} - \frac{\xi^4}{6R^4}\right) \ , \tag{S.10}$$

where $E$ is the elastic modulus of the surrounding elastic network and $c^0_{\text{out}}$ is the equilibrium concentration outside the condensate in thermodynamic limit (i.e., large condensate size leading to negligible curvature). This particular form of mechanical pressure is obtained by considering the expansion of a pore (i.e., the droplet in our case) in a neo-Hookean material [1].

Furthermore, the conservation of mass implies

$$\bar{c}V = c_{\text{in}} \frac{4\pi R^3}{3} + c_\infty \left(V - \frac{4\pi R^3}{3}\right) \tag{S.11}$$

where $\bar{c}$ is the average concentration and $V$ is the system volume (i.e., the volume of the nucleus). Assuming $V \gg 4\pi R^3/3$,

$$c_\infty \approx \bar{c} - c_{\text{in}} \frac{4\pi R^3}{3V} \ . \tag{S.12}$$

The resultant equation for droplet growth, derived using Eq. S.12, Eq. S.9 and Eq. S.7, is given by

$$\frac{dR}{dt} = \frac{D}{Rc_{\text{in}}} \left(\bar{c} - c_{\text{in}} \frac{4\pi R^3}{3V} - c^0_{\text{out}} \exp\left(\frac{P}{c_{\text{in}} k_B T}\right)\right) \ , \tag{S.13}$$

which is the same as the dynamical equation of condensate growth presented in the main text. The above equation can be re-written as:

$$\frac{dR}{dt} = \left[\frac{a}{R} - bR^2 - \frac{c}{R}\exp(\frac{2\tilde{\gamma}}{R} + \tilde{E}g(R))\right] , \tag{S.14}$$

with parameters: $a = D\bar{c}/c_{\text{in}}$, $b = 4\pi D/3V$, $c = Dc^0_{\text{out}}/c_{\text{in}}$, $\tilde{\gamma} = \gamma/(c_{\text{in}}k_BT)$, $\tilde{E} = E/(c_{\text{in}}k_BT)$ and $g(R) = \left(\frac{5}{6} - \frac{2\xi}{3R} - \frac{\xi^4}{6R^4}\right)$. This form of the equation is used for fitting model to experimental data.

##### B. Growth of multiple condensates

We extend the description of a single condensate growth to the case of N growing condensates/droplets in an elastic network surrounding them. Material flux balance, as introduced before in Eq. S.7, becomes

$$\frac{dR_i}{dt} = \frac{D}{c_{\text{in}}R_i}\left(c_\infty - c_{\text{out (i)}}\right) \tag{S.15}$$

where $R_i$ is the radius of the $i^{th}$ droplet and $c_{\text{out (i)}}$ is the concentration of the dilute phase outside the $i^{th}$ droplet. The concentration of the dilute phase just outside the droplets can be different due to the difference in local mechanical properties. We have assumed the diffusion of the dilute phase to be the same everywhere and $c_{\text{in}}$ to be the same for all droplets. Material conservation for $N$ droplets leads to the equation

$$c_\infty(V - \sum_{i=1}^{N} V_i^d) + c_{\text{in}}\sum_{i=1}^{N} V_i^d = \bar{c}V \tag{S.16}$$

where $V_i^d$ is the volume of the $i^{th}$ droplet and $V$ is the system volume. Using the above Eq. S.15 & S.16 we can arrive at the dynamical equation for the growth of the $i^{th}$ droplet:

$$\frac{dR_i}{dt} = \frac{D}{c_{in}R_i}\left(\bar{c} - c_{in}\sum_{j}^{N}\left(\frac{4\pi R_j^3}{3V}\right) - c^0_{out}\exp(\frac{P_i(R_i)}{c_{in}K_BT})\right) , \tag{S.17}$$

where pressure on the $i^{th}$ condensate surface is given by

$$P(R_i) = \frac{2\gamma}{R_i} + E_i\left(\frac{5}{6} - \frac{2\xi_i}{3R_i} - \frac{\xi_i^4}{6R_i^4}\right) \tag{S.18}$$

here $\xi_i$ and $E_i$ are the local mesh size and local stiffness of the network around the $i^{th}$ droplet. Using the parameters defined in the previous section we arrive at the dynamical equation

$$\frac{dR_i}{dt} = \frac{a}{R_i} - \frac{b}{R_i}\sum_{j=1}^{N} R_j^3 - \frac{c}{R_i}\exp\left(\frac{2\tilde{\gamma}}{R_i} + \tilde{E}_i(\frac{5}{6} - \frac{2\xi_i}{3R_i} - \frac{\xi_i^4}{6R_i^4})\right) . \tag{S.19}$$

##### C. Mean field theory of condensate coarsening

Growth and ripening of condensates embedded in an elastic media can be investigated using the classical mean-field approximation developed by Lifshitz, Slyozov and Wagner [2, 3]. Let us first revisit the case of a fluid droplet growing in the absence of an elastic medium. We can rewrite Eq. S.15 using Eq. S.9 as

$$\frac{dR}{dt} = \frac{D}{c_{\text{in}} R}\left(c_\infty - c_{\text{out}}^0\left(1 + \frac{2\gamma}{c_{\text{in}} K_B T R}\right)\right) \;, \tag{S.20}$$

where we have taken a linear approximation of the exponential term. The above equation can be rewritten as

$$\frac{dR}{dt} = \frac{D c_{\text{out}}^0}{R c_{in}}\left(\Delta - \frac{l_c}{R}\right) \tag{S.21}$$

where $l_c = \frac{2\gamma}{c_{\text{in}} K_B T}$ is the capillary length and $\Delta = \frac{c_\infty - c_{\text{out}}^0}{c_{\text{out}}^0}$ is the extent of supersaturation of the condensate proteins in the medium. The above equation (Eq. S.21) dictates the stability of the droplets with an unstable fixed point in dynamics of $R$ at critical radius $R_c = l_c/\Delta$. Droplets with size below the critical radius would shrink, whereas those larger than $R_c$ would grow. Note that both $\Delta$ and $R_c$ are functions of time. In the classical LSW theory, Eq. S.21 was considered as a mean-field description of condensates going through ripening to find the asymptotic scaling law of mean condensate size to be $\langle R\rangle \sim t^{1/3}$.

Next, we consider the growth of a droplet in the presence of a homogeneous elastic network with stiffness $E$. Including the elastic pressure term in Eq. S.20 we obtain:

$$\begin{aligned}\frac{dR}{dt} &= \frac{D c_{\text{out}}^0}{R c_{\text{in}}}\left[\Delta - \left(\frac{l_c}{R} + \tilde{E}\left(\frac{5}{6} - \frac{2\xi}{3R} - \frac{\xi^4}{6R^4}\right)\right)\right] \\ &\simeq \frac{D c_{\text{out}}^0}{R c_{\text{in}}}\left[\left(\Delta - \frac{5\tilde{E}}{6}\right) - \frac{1}{R}\left(l_c - \frac{2\xi\tilde{E}}{3}\right)\right] \\ &= \frac{D c_{\text{out}}^0}{R c_{\text{in}}}\left(\tilde{\Delta} - \frac{\tilde{l}_c}{R}\right)\end{aligned} \tag{S.22}$$

where the $\tilde{\Delta} = \Delta - \frac{5\tilde{E}}{6}$ is the renormalized measure of supersaturation and $\tilde{l}_c = l_c - \frac{2\xi\tilde{E}}{3}$ is the renormalized capillary length. Note that unlike in the LSW description, the renormalized supersaturation and capillary length can be negative depending on the stiffness in the embedding network. In the case where both $\tilde{\Delta}, \tilde{l}_c > 0$, the system will retain the unstable fixed point and the system will go through Ostwald ripening. In the case $\tilde{\Delta} < 0$, the droplets shrink at any size due to elasticity-driven suppression of phase separation. In the other regime where $\tilde{\Delta} > 0$, $\tilde{l}_c < 0$, droplets of any size will grow (a stable fixed point at $\infty$) until the growth is stopped by lack of supersaturation $\Delta \to 0$. This corresponds to the suppression of Ostwald ripening due to the effect of the surrounding elastic network. Elastic ripening is not possible in this case as we have assumed a homogeneous elastic medium.

In the case of a heterogeneous medium where droplets grow at varying stiffness values, both the degree of supersaturation ($\tilde{\Delta}$) and the capillary length ($\tilde{l}_c$) can vary among droplets. The mean-field approximation employed in the LSW approach becomes inapplicable under these conditions. Consequently, we have devised an approximate linear theory based on the size difference between droplets ($\Delta R$) to explore the various forms of ripening and suppressed ripening in condensates within heterogeneous elastic networks.

#### II. LINEARIZED THEORY OF DROPLET RIPENING

Here we derive an approximate theory of ripening in terms of difference in condensate size $\Delta R = R_2 - R_1$ to understand the three possible scenarios of condensate coarsening – (i) Ostwald ripening, (ii) suppressed ripening, and (iii) elastic ripening. We rewrite the equation for the size dynamics of the $i^{th}$ droplet Eq. S.19 as:

$$\frac{1}{2}\frac{dR_i^2}{dt} \;=\; a - b\sum_j R_j^3 - c\exp\left(\frac{2\tilde{\gamma}}{R_i} + \tilde{E}_i(\frac{5}{6} - \frac{2\xi}{3R_i} - \frac{\xi^4}{6R_i^4})\right) \tag{S.23}$$

Now, for two droplets of radii $R_1$ and $R_2$, we use Eq. S.23 to arrive at

$$\frac{1}{2}\frac{d}{dt}\left(R_1^2 - R_2^2\right) \;=\; -c\exp\left(\frac{2\tilde{\gamma}}{R_1} + \tilde{E}_1(\frac{5}{6} - \frac{2\xi}{3R_1} - \frac{\xi^4}{6R_1^4})\right) + c\exp\left(\frac{2\tilde{\gamma}}{R_2} + \tilde{E}_2(\frac{5}{6} - \frac{2\xi}{3R_2} - \frac{\xi^4}{6R_2^4})\right), \tag{S.24}$$

and by taking a linear approximation of the exponential term we get

$$\frac{1}{2}\frac{d}{dt}\left((R_1+R_2)(R_1-R_2)\right) \;=\; -c\left(2\tilde{\gamma}(\frac{1}{R_1} - \frac{1}{R_2}) + \frac{5\Delta\tilde{E}}{6} - \frac{2\xi}{3}\left(\frac{\tilde{E}_1}{R_1} - \frac{\tilde{E}_2}{R_2}\right) - \frac{\xi^4}{6}\left(\frac{\tilde{E}_1}{R_1^4} - \frac{\tilde{E}_2}{R_2^4}\right)\right). \tag{S.25}$$

We consider a case where the two droplets have grown after nucleation and have reached a similar size $R_1 \sim R_2 \sim R$ but there is a stiffness difference between them $\tilde{E}_1 > \tilde{E}_2$. The droplets start ripening with a small size difference $\Delta R = R_2 - R_1$, with $R_2 > R_1$. We also assume that the overall growth rate of the two droplets to be very slow $\frac{d}{dt}\left((R_1 + R_2)\right) \sim 0$, owing to volume conservation and $\Delta R \ll R$. Using these considerations and ignoring higher order terms in $\frac{\xi}{R}$, we can write the equation for $\Delta R$ as

$$R\frac{d\Delta R}{dt} = c\left(\left(2\tilde{\gamma} - \frac{2\xi}{3}\tilde{E}_1\right)\frac{\Delta R}{R^2} + \left(\frac{5}{6} - \frac{2\xi}{3R}\right)\Delta\tilde{E}\right) \tag{S.26}$$

This can be rewritten as

$$\frac{d\Delta R}{dt} = \frac{c}{R}\left(\left(2\tilde{\gamma} - \frac{2\xi}{3}\langle\tilde{E}\rangle\right)\frac{\Delta R}{R^2} + \left(\frac{5}{6} - \frac{2\xi}{3R}\right)\Delta\tilde{E}\right), \tag{S.27}$$

which we can identify as the equation for $\Delta R$ dynamics presented in the main text.

#### III. DROPLET NUCLEATION AND GROWTH IN HETEROGENEOUS ELASTIC LANDSCAPE

According to the classical nucleation theory, nucleation of a droplet depends on the competing effects of condensation into the fluid phase and the energetic cost of surface tension at the interface between the condensed and the dilute phases. In the presence of an elastic network, the nucleated condensates must also deform the surrounding network to expand, thus incurring an additional elastic energy cost. The change in Gibbs free energy to nucleate a droplet of radius $R$ in a network of stiffness $E$ is given by

$$\Delta G = 4\pi R^2 \gamma - \frac{4}{3}\pi R^3 \left(c_{\text{in}} k_B T \log\left(S\right) - P_E\right) , \tag{S.28}$$

where $P_E$ is the pressure due to elastic deformation of the network and $S$ is the extent of super-saturation given by $S = \frac{P(T)}{P_{\text{eq}}(T)} \sim \frac{c_{\text{out}} V}{c^* V}$. Here $P$ and $P_{\text{eq}}$ are the pressures of the supersaturated protein solution and the equilibrium pressure at that temperature. The parameters $c_{\text{in}}$, $c_{\text{out}}$ and $c^*$ are the concentration of molecules in the condensed phase, dilute phase and the critical density at equilibrium ($S = 1$). The extent of supersaturation decreases as more and more droplets are nucleated. Using the conservation of droplet material $\bar{c}V = c_{\text{in}} V_R + c_{\text{out}}(V - V_R)$, we can write $c_{\text{out}} = \bar{c} - c_{\text{in}} \frac{V_R}{V}$, where $V_R = \sum_i^N \frac{4}{3}\pi R_i^3$ is the instantaneous total droplet volume and $V$ is the system size. Thus, as the droplets are nucleated the supersaturation $S = c_{\text{out}}/c^*$ would decrease. While droplet growth can also reduce supersaturation, we assume that the timescale of nucleation is much faster than the timescale of growth such that nucleation is complete before the droplets can grow significantly [4]. Using the expression for pressure (Eq. S.10) due to the elastic deformation of the network we can write

$$\frac{\Delta G}{c_{\text{in}} k_B T} = 4\pi R^2 \frac{\gamma}{c_{\text{in}} k_B T} - \frac{4}{3}\pi R^3 \left(\log\left(\frac{\bar{c}}{c^*} - \frac{c_{\text{in}}}{c^*}\frac{V_R}{V}\right) - \frac{E}{c_{\text{in}} k_B T} g(R)\right) . \tag{S.29}$$

where $g(R) = \left(\frac{5}{6} - \frac{2\xi}{3R} - \frac{\xi^4}{6R^4}\right)$. We can rewrite the above equation with our previously defined scaled parameters as

$$\frac{\Delta G}{c_{\text{in}} k_B T} = 4\pi R^2 \tilde{\gamma} - \frac{4}{3}\pi R^3 \left(\log\left(\frac{\bar{c}}{c^*} - \frac{c_{\text{in}}}{c^*}\frac{V_R}{V}\right) - \tilde{E} g(R)\right) . \tag{S.30}$$

The probability of nucleating a droplet of size $R$ is given by $p_{\text{nuc}}(R, E) = (1/Z)\, e^{-\frac{\Delta G(R,E)}{k_B T}}$ where $Z = \int_0^\infty e^{-\frac{\Delta G(R)}{k_B T}}\, dR$ is the normalization constant. This result can be used to stochastically simulate the nucleation of droplets. We consider a heterogeneous elastic network with the normal distribution of stiffness values with a given mean stiffness $\langle E \rangle$ and a coefficient of variation $CV_E$. To nucleate the $i^{th}$ droplet we pick a nucleation size $R_i^0$ from a normal distribution, draw a stiffness value $E_i$ from the stiffness distribution, and then calculate the probability of nucleation of a droplet of size $R_i$ at stiffness $E_i$ by $p_{\text{nuc}}(R_i^0, E_i)$.

The local mesh size $\xi_i$ was calculated as $\xi_i \sim E_i^{-1/4}$, based on the relation of stiffness and mesh size with local density ($\rho$) given by $\sqrt{E_i} \sim \rho$ and $\xi_i^{-2} \sim \rho$ [5, 6]. This process of nucleation stops when supersaturation $S$ reaches 1 or when a maximum number of droplets have nucleated or after a maximum number of tries ($\sim$ time duration of the nucleation period). The constraints on the droplet number and nucleation period were tuned to match the experimental results. This nucleation dynamics leads to experimentally observed features of condensate growth inside the nucleus such as the condensates predominantly nucleate in the softer regions (i.e., low density) of the chromatin network [7] (see Fig. S3).

To study how the stiffness of the chromatin network affects the condensate growth, we simulate condensate nucleation and growth in a varying stiffness landscape as dicussed above (Fig. S4A). We calculate the ripening fraction (i.e., the fraction of condensates that shrank to zero during ripening) and the scaling coefficients during growth with varying the mean and coefficient of variation of the stiffness distribution (Fig. S4B,C,D). The treatment with TSA reduces chromatin heterogeneity (see main text Fig. 4) and makes the chromatin softer. This reduction in stiffness of the surrounding elastic network will decrease the equilibrium concentration of the dilute phase $c_{\text{out}}$ (see Eq. S.9), which in turn will increase the level of supersaturation given the same amount of overall protein concentration ($\bar{c}$). Thus, we model TSA-treated chromatin network as a stiffness landscape with lower values of mean stiffness $\langle E \rangle$, a lower coefficient of variation of stiffness $CV_E$ and a higher value of initial supersaturation (Fig. S4C,D).

**SUPPLEMENTARY FIGURES**

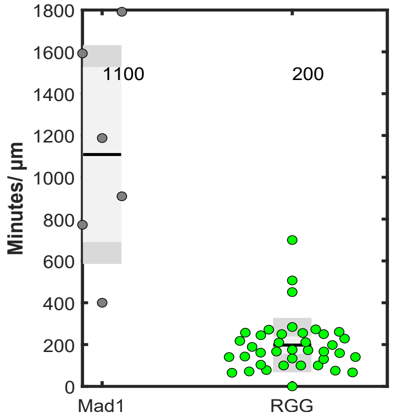


**Figure S** 1. Condensate ripening time per micron for the coiled-coil condensates and the disordered protein condensates.

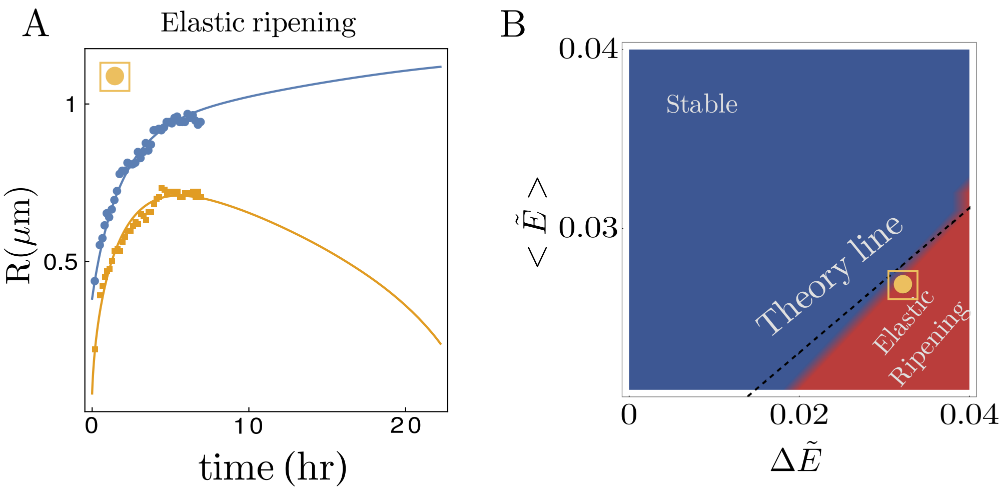


**Figure S 2.** Elastic ripening in coiled-coil condensates. (A) Theoretical fit (lines) to the experimental data (points) indicate ripening of the smaller droplet growing in a stiffer chromatin environment. (B) A theoretical phase diagram in the plane of mean stiffness ($\langle\tilde{E}\rangle$) and the difference in stiffness ($\Delta\tilde{E}$) shows suppressed ripening and elastic ripening behaviours. The dashed line is a prediction from the linear theory of ripening. The yellow point denotes the position of the experimentally observed data estimated by the fitting shown in panel A. Model parameter values are: $D\bar{c}/c_{\text{in}} = 0.0003\ \mu m^2 s^{-1}$, $4\pi D/3V = 0.00003\ \mu m^{-1} s^{-1}$, $Dc^0_{\text{out}}/c_{\text{in}} = 0.00026$, $\tilde{\gamma} = 0.0005\,\mu m$, $\tilde{E}_1 = 0.010750$, $\tilde{E}_2 = 0.042989$.

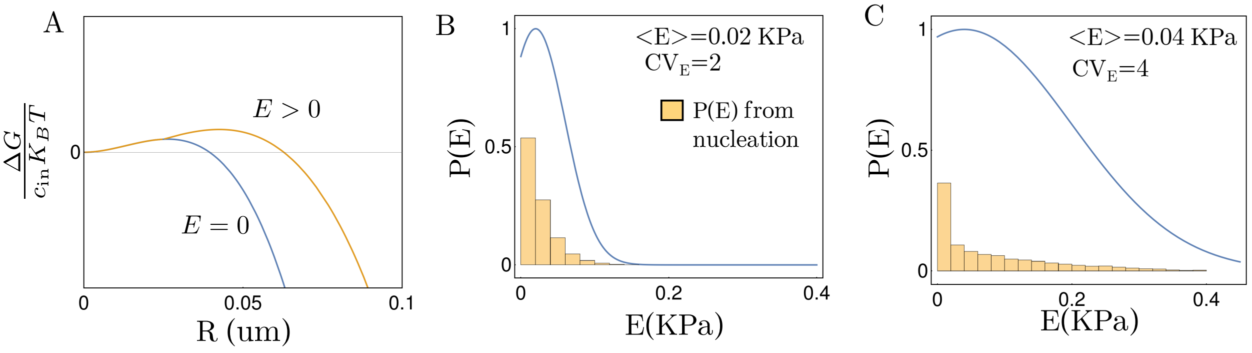


**Figure S 3.** Nucleation of condensates in heterogeneous elastic environment. (A) Energy cost of nucleation as a function of condensate size $R$. (B& C) Stiffness distribution of the heterogeneous media (lines) and histogram of stiffness where droplets are nucleated (bar plots). The parameter values are the same as used in Fig 3F in the main text.

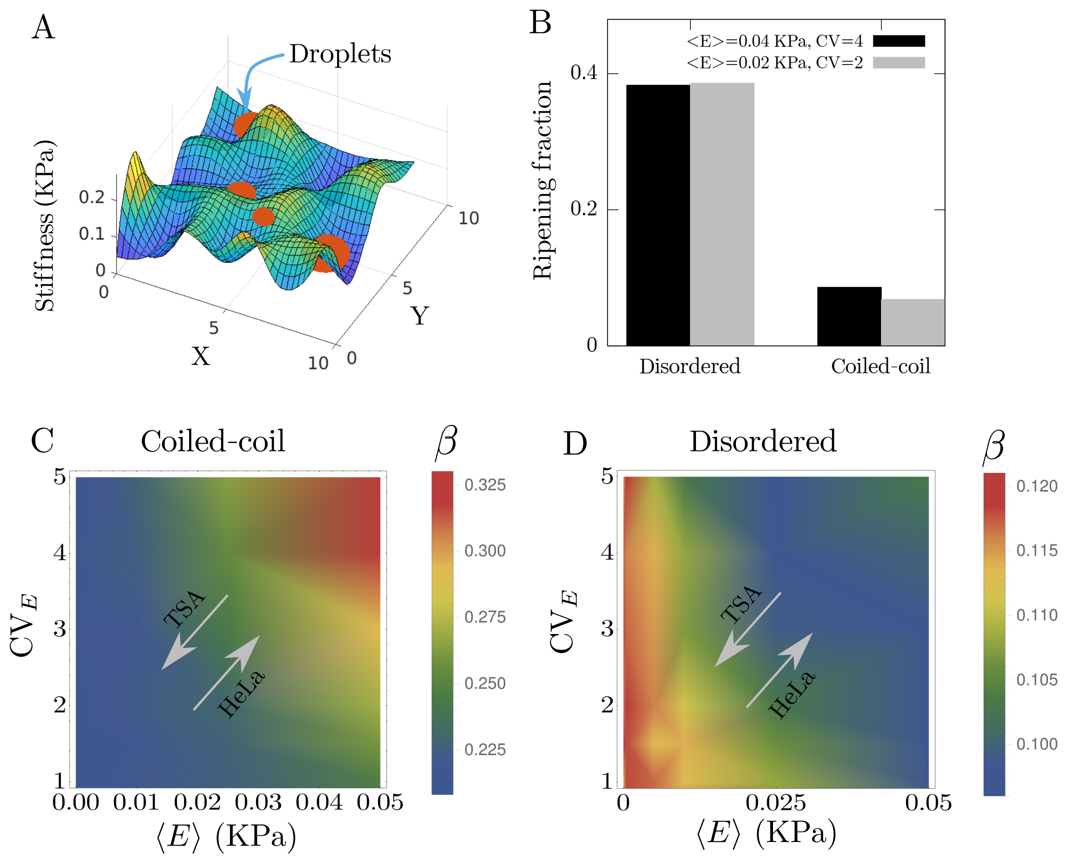


**Figure S** 4. Condensate size scaling. (A) A snapshot of condensates growing in a heterogeneous elastic medium. (B) Average ripening fraction for disordered and coiled-coil condensates. The fractions do not change significantly with changing the mean and the coefficient of variation of the stiffness distribution. (C) Phase diagram of the scaling coefficient $\beta$ with changing mean stiffness ($\langle E \rangle$) and coefficient of variation of stiffness ($\mathrm{CV}_E$) for coiled-coil protein condensates. (D) Phase diagram of the scaling coefficient $\beta$ with changing mean stiffness ($\langle E \rangle$) and coefficient of variation of stiffness ($\mathrm{CV}_E$) for disordered protein condensates. The parameter values are the same as used in Fig 3F in the main text.

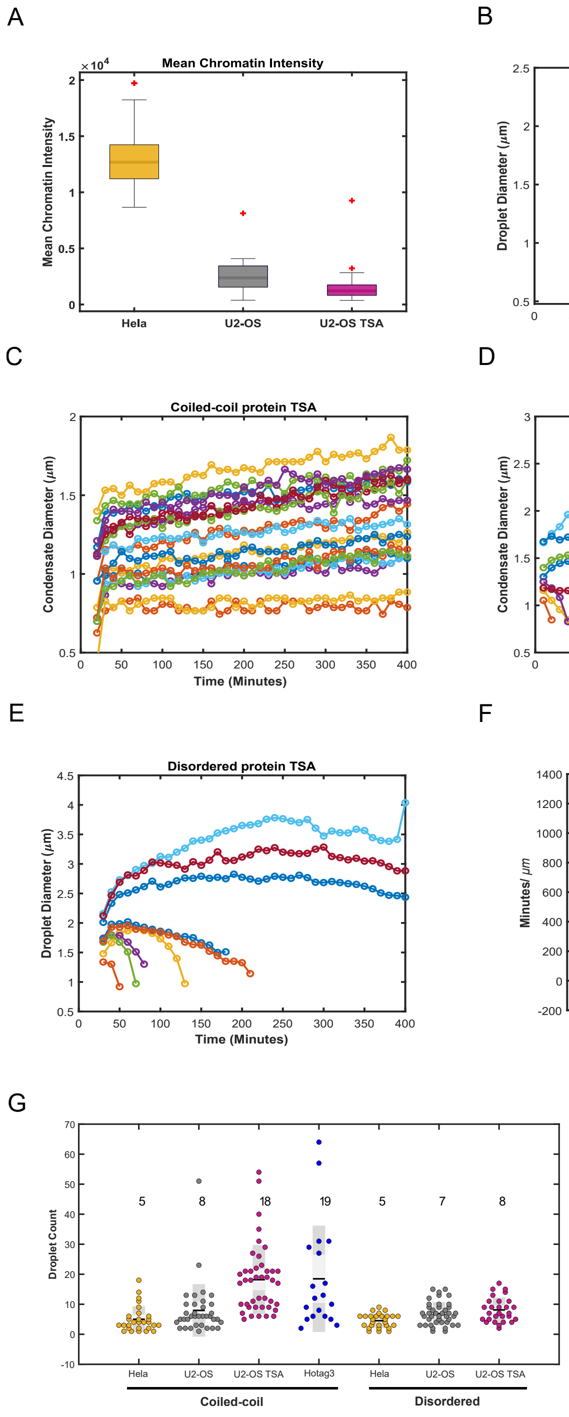


**Figure S 5.** Effect of chromatin environment on condensate growth dynamics. (A) Box plot of the mean chromatin intensity of HeLa, U2-OS and TSA treated U2-OS cells. (B) Representative growth patterns of coiled-coil condensates in HeLa nuclei. (C) Representative growth patterns of coiled-coil condensates in TSA treated U2-OS nuclei. (D) Representative growth patterns of disordered condensates in HeLa nuclei. (E) Representative growth patterns of disordered condensates in TSA treated U2-OS nuclei. (F) The ripening times per micrometer for the condensates in HeLa, U2-OS and in untreated U2-OS nuclei (averages listed on plot). (G) The condensate count per cell for the proteins in different chromatin environment (averages listed on plot).
